# S-Acylation Regulates the Membrane Association and Activity of Calpain-5

**DOI:** 10.1101/2022.01.25.477766

**Authors:** Jozsef Gal, Vimala Bondada, Charles B. Mashburn, David W. Rodgers, Dorothy E. Croall, James W. Geddes

## Abstract

Calpain-5 (CAPN5) is a member of the calpain family of calcium-activated neutral thiol proteases. CAPN5 is partly membrane associated, despite its lack of a transmembrane domain. Unlike classical calpains, CAPN5 contains a C-terminal C2 domain. C2 domains often have affinity to lipids, mediating membrane association. We recently reported that the C2 domain of CAPN5 was essential for its membrane association and the activation of its autolytic activity. However, despite the removal of the C2 domain by autolysis, the N-terminal fragment of CAPN5 remained membrane associated. S-acylation, also referred to as S-palmitoylation, is a reversible post-translational lipid modification of cysteine residues that promotes membrane association of soluble proteins. In the present study several S-acylated cysteine residues were identified in CAPN5 with the acyl-PEG exchange method. Data reported here demonstrate that CAPN5 is S-acylated on up to three cysteine residues including Cys-4 and Cys-512, and likely Cys-507. The D589N mutation in a potential calcium binding loop within the C2 domain interfered with the S-acylation of CAPN5, likely preventing initial membrane association. Mutating specific cysteine residues of CAPN5 interfered with both its membrane association and the activation of CAPN5 autolysis. Taken together, our results suggest that the S-acylation of CAPN5 is critical for its membrane localization which appears to favor its enzymatic activity.

## Introduction

Calpains are calcium-dependent, non-lysosomal thiol proteases that regulate biological processes by either targeting substrate proteins for degradation or modifying the activity of the proteolytic products [1, 2]. Calpains are defined by the presence of the calpain catalytic domain that is made up by two subdomains. There are 15 calpains encoded by the human genome. Some are expressed ubiquitously, whereas others show distinct developmental and/or tissue-specific expression patterns [3, 4]. In the catalytic center of calpains, three amino acid residues: a cysteine, a histidine and an asparagine are critical to carry out proteolysis of the substrate proteins [5].

Historically, much of the research focused on classical calpains 1 and 2, both of which are ubiquitous and heterodimeric. They are designated classical calpains based on the presence of a C-terminal penta-EF hand (PEF) domain in both the catalytic (CAPN1 or CAPN2) and small subunit (CAPNS1 formerly called CAPN4) [4]. The PEF domains bind Ca^2+^ and mediate the interaction between the two subunits. Non-classical calpains lack the C-terminal PEF domain [6] and it is suggested that these calpains are evolutionarily older [7, 8]. Within this group, calpain-5 (CAPN5) and calpain-6 (CAPN6) have C-terminal domains designated as C2 domains which are often involved in Ca^2+^-dependent lipid interactions and membrane localization [9]. CAPN6, an isoform that lacks the active site cysteine, was reported to be microtubule associated [10] with expression limited to placenta, early development, and regenerating muscle [11–13]. In contrast to the restricted expression of CAPN6, CAPN5 is ubiquitously expressed at all stages of development [12].

The discovery that mutations in CAPN5 are causal for a rare devastating eye disease [14–17] brought increased attention to this non-classical calpain. CAPN5 was formerly called hTra3 after its orthologue in *C. elegans* [18]. CAPN5 is relatively understudied although its expression at the mRNA level was the second highest among calpains in the rat brain [19]. Subcellular localization of the CAPN5 enzyme is expected to provide clues as to its relevant substrates and physiological function(s) but is not yet fully understood. CAPN5 is detected in a variety of subcellular fractions including nuclear, cytosolic, ER, and a crude synaptic-mitochondrial fraction, but results were influenced greatly by which antibody was used [20].

Immunohistochemistry has suggested that CAPN5 is predominantly localized near the cell surface in a ruffled pattern [14], although cytosolic and nuclear localization have also been observed [2, 19]. Proteomic studies of subcellular fractions from hepatocytes and erythrocytes also document the presence of endogenous CAPN5 within the membrane fraction [21, 22].

The mechanism of the association of CAPN5 with the plasma membrane and membrane-bound organelles, despite the absence of any predicted transmembrane domain in the protein, is not well understood [14, 21, 22]. CAPN5 harbors a predicted C-terminal C2 domain. C2 domains are often mediators of membrane or lipid binding and are frequently also calcium regulated [9], although some C2 domains are non-Ca^2+^ binding and some are not associated with enzymatic activity such as that of CAPN6. We recently demonstrated that the CAPN5 C2 domain has a functional role in mediating membrane association of CAPN5 in SH-SY5Y cells [2].

In the presence of added Ca^2+^, membrane-associated CAPN5 undergoes autolysis [23], resulting in removal of a substantial portion of the C2 domain [2]. However, the N-terminal fragment that contains the catalytic core of CAPN5 remains associated with membrane, despite the loss of the C2 domain. This suggests that CAPN5 has another mode of membrane association or anchoring. We hypothesized that in addition to the C2 domain, the membrane association of CAPN5 may also be mediated by S-acylation. This was based on the observation that approximately 18% of proteins encoded by the human genome may be S-acylated [24], and that in subcellular fractionation studies the apparent molecular weight of full-length CAPN5 differed among antibodies [20]. Protein S-acylation can result in a mobility shift [25]

Protein S-acylation is the covalent attachment of long-chain fatty acids, either saturated or unsaturated, to cysteine residues through thioester linkage [26, 27]. Because the most abundantly attached fatty acid is the 16-carbon saturated palmitate (C16:0), protein S-acylation is also commonly referred to as S-palmitoylation. However, stearate (C18:0), oleate (C18:1), and other long-chain fatty acids are also attached through thioester bond [28–30]. S-acylation supports or modulates the membrane association of transmembrane proteins and peripheral membrane proteins anchored via other lipid modifications such as myristoylation [31]. However, unlike other protein lipid modifications, S-acylation is reversible. Proteins can undergo cycles of S-acylation, mediated by protein S-acyltransferases (PATs) and de-acylation via acyl-protein thioesterases (APTs) [26, 32].

The aim of this study was to evaluate the S-acylation of human CAPN5 in the SH-SY5Y neuroblastoma cell line by using an acyl-polyethylene glycol (PEG) exchange (APEx) based method [33]. The advantage of this strategy is that information about the number of S-acylated sites may be determined with analysis of the APEx treated samples by denaturing gel electrophoresis and immunoblotting. Data reported here demonstrate that over half (59%) of exogenously expressed CAPN5 is S-acylated on one, two, or three cysteine residues including Cys-4 and Cys-512, and likely Cys-507. Alteration of a potential calcium binding loop within the C2 domain negatively affected acylation of CAPN5, suggesting a model where the C2 domain has a role in facilitating acylation by localizing CAPN5 with the enzymes required for S-acylation. Additionally, immunohistochemistry and subcellular fraction demonstrate the importance of S-acylation for membrane association of CAPN5 and retention of a catalytic fragment of CAPN5 at the membrane.

## Materials and Methods

### Plasmid construction

The generation of the C-terminally 3×FLAG-tagged wild-type (WT), C81A and D589N mutant CAPN5 expression constructs (phCAPN5-3×FLAG) and the corresponding stable cell lines were reported elsewhere [2]. The C4A mutation was introduced by PCR amplification of the CAPN5 cDNA using an upstream primer that contained the mutation and inserting the amplification product between the *Eco*RI and *Xba*I sites of p3×FLAG-CMV-14 (Sigma, E4901). The C20A, C377A, C495A, and C507A CAPN5 mutations and the C506A CAPN6 mutation were introduced with the QuikChange II Site-Directed Mutagenesis Kit (Agilent, 200523). The C246A and C512A mutations and the C495A/C507A/C512A triple mutation were introduced by PCR amplification of phCAPN5-3×FLAG with 5’-phosphorylated primers incorporating the respective mutations, followed by circularization with ligation.

A 58 bp deletion [34] was introduced into pcDNA3.1/Hygro(+) (Thermo Fisher Scientific, V87020) between the *Bpl*I and *Xma*I sites to inactivate the SV40 origin of replication without inactivating the SV40 promoter that drives the expression of the hygromycin resistance gene. This hygromycin resistance cassette was isolated as a *Csi*I-*Bsm*I fragment, and used to replace the corresponding fragment of phCAPN5-3×FLAG to generate a construct that is selectable by hygromycin in mammalian cells. The C-terminally 3×HA-tagged CAPN5 expression construct was made by replacing the *Bam*HI-*Xma*I small fragment of phCAPN5-3×FLAG encoding the 3×FLAG tag with annealed oligonucleotides.

The C-terminally 3×FLAG-tagged human CAPN6 expression construct was made by PCR amplification of the CAPN6 coding region from MGC clone 3503916 as template (Dharmacon, MHS6278-202828128), and inserting it between the *Eco*RI and *Xba*I sites of p3×FLAG-CMV-14. The p3×FLAG-CAPN1 and p3×FLAG-CAPN2 expression constructs were gifts from Dr. Yi Zhang (Addgene plasmids # 60941 and 60942, http://n2t.net/addgene:60941, RRID:Addgene_60941, and http://n2t.net/addgene:60942, RRID:Addgene_60942 [35]). The DNA oligonucleotides used in this work are summarized in Table 1. All constructs were confirmed with sequencing.

**Table 1.**
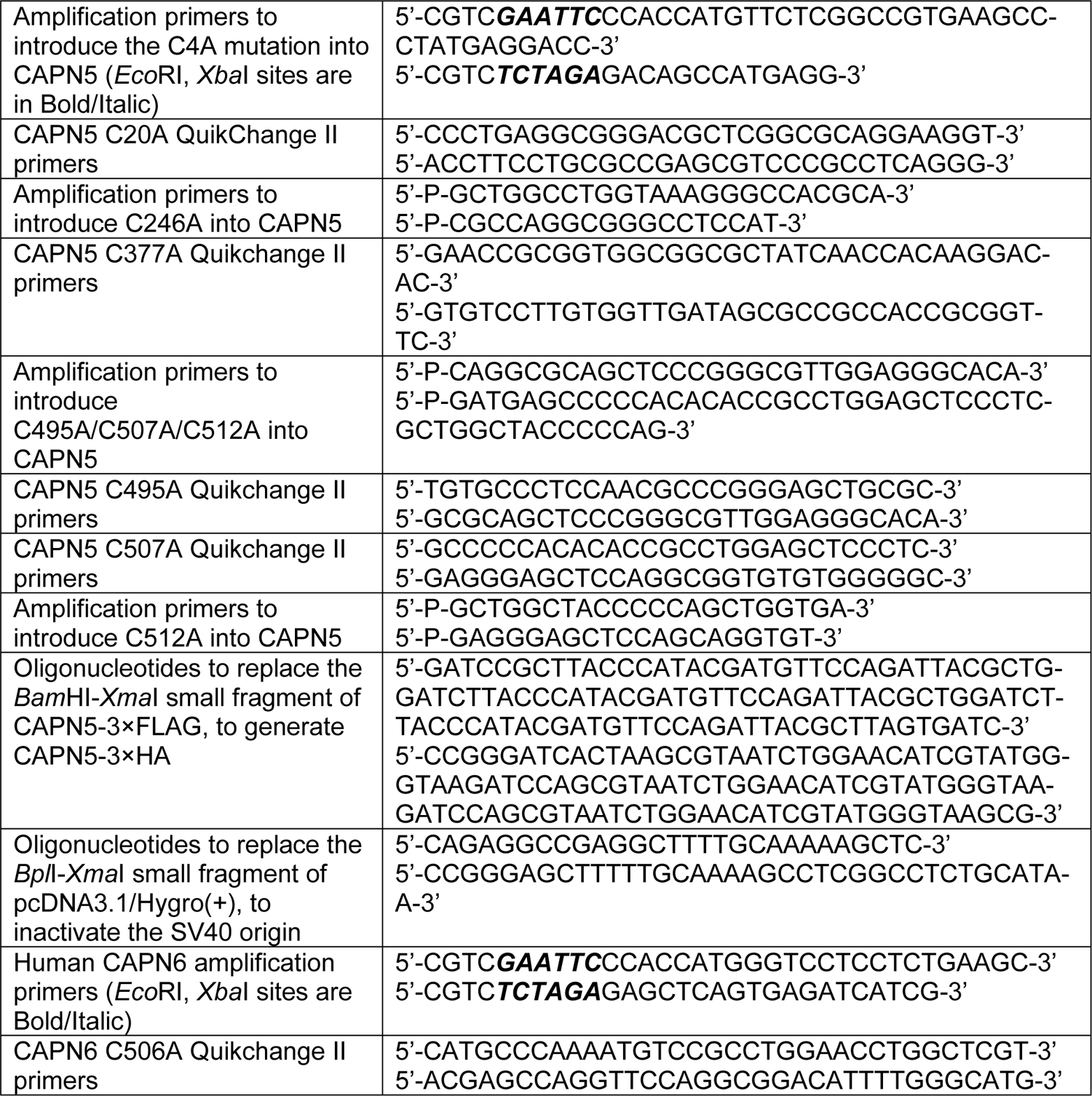
The DNA oligonucleotides used in this study. P, phosphate.

### Cell culture and the generation of stable cell lines

The human neuroblastoma cell line SH-SY5Y (ATCC, CRL-2266) and its derivatives were maintained in MEM with Earle’s salts and L-glutamine (Corning, 10-010-CV) supplemented with 10% fetal bovine serum (FBS) (Atlanta Biologicals, S11195H) and penicillin-streptomycin (Corning, 30-002-CI). The 293T cells (ATCC, CRL-3216) [36] were cultured in high glucose DMEM (Sigma, D5796) supplemented with 10% FBS and penicillin-streptomycin. The cells were maintained in humidified cell culture incubator under 5% CO_2_/95% air at 37°C. The cells were transfected with the respective expression constructs using the LipoJet DNA In Vitro Transfection Reagent (SignaGen, SL100468). Stable cells were selected and maintained with 2,500 µg/ml G-418 (Gold Biotechnology, G-418-25) in the medium with the exception of the 293T-based CAPN5-3×FLAG stable cell line that was selected and maintained with 200 µg/ml hygromycin (Gold Biotechnology, H-270-1).

### Acyl-PEG exchange (APEx)

The acyl-PEG exchange experiments [33] were performed with the SiteCounter S-Palmitoylated Protein Kit (Badrilla, K010-312), following the manufacturer’s instructions. For each reaction, half of the cells from a confluent 10 cm culture dish were used. Briefly, the cells were lysed, and the free thiol groups were blocked. The thioester bonds were cleaved by hydroxylamine treatment, followed by tagging of the acyl-free cysteine residues with polyethylene glycol (PEG) groups. Negative control samples were also prepared without thioester cleavage as a control for the initial thiol blocking step. A small modification from the kit protocol was that after cell lysis, the samples were passed through 18-gauge, then 23-gauge needles several times before the acetone precipitation step to decrease their viscosity.

### Subcellular fractionation

The cytosolic and membrane fractions were prepared using the Mem-PER Plus Membrane Protein Extraction Kit (Thermo Fisher Scientific, 89842) with slight modifications. The samples were kept on ice unless otherwise indicated, and all centrifugations were performed at 4°C. Briefly, the cells were grown to 100% confluence, trypsinized, and pelleted by centrifugation at 1,200 × g for 10’. The trypsinized cells were washed two times with Cell Wash Solution, followed by centrifugation at 500 × g for 6’. The cell pellets were resuspended in Permeabilization Buffer supplemented with 2 mM calcium chloride and protease inhibitors (cOmplete Mini EDTA-free Protease Inhibitor Cocktail, Sigma, 11836170001), vortexed briefly, and incubated for 30’ at 4°C with rotation. After incubation, the permeabilized cells were centrifuged at 16,000 × g for 15’. The supernatant (cytosolic fraction) was carefully collected. To eliminate cytosol carryover, the pellets were incubated with cold 0.1 M PBS (72 mM Na_2_HPO_4_, 28 mM NaH_2_PO_4_, 0.9% NaCl, pH 7.4) supplemented with protease inhibitors for 5’ at 4°C with rotation, followed by centrifugation at 16,000 × g for 10’. The washed pellets were incubated in Solubilization Buffer supplemented with protease inhibitors for 30’ and centrifuged at 16,000 × g for 15’, and the supernatant (membrane fraction) was collected. Protein concentrations were determined with BCA protein assay (Thermo Fisher Scientific, 23225), and equal protein amounts were used in denaturing gel electrophoresis.

### Denaturing gel electrophoresis and immunoblotting

Following the APEx experiments, equal volumes of the total extracts (“input”), the PEG-tagged samples and the negative controls of the respective stable cell lines were resolved by denaturing gel electrophoresis on 3-8% Tris-acetate gradient protein gels (Thermo Fisher Scientific, EA0378BOX) using Tris-acetate-SDS running buffer, pH 8.24 (Thermo Fisher Scientific, LA0041). The level of C81A mutant CAPN5 was higher than that of WT CAPN5, whereas all other CAPN5 mutants were expressed at lower levels (Supplemental Figure 1). Accordingly, the loading of the APEx samples was adjusted to obtain comparable anti-FLAG signal in immunoblotting. The subcellular fractionation samples were resolved by denaturing gel electrophoresis on 4-12% Bis-Tris gradient protein gels (Thermo Fisher Scientific, NP0335BOX) using MES SDS running buffer, pH 7.3 (Thermo Fisher Scientific, NP0002). The resolved proteins were transferred to nitrocellulose membranes with 0.2 µm pore size (Bio-Rad, 1704158), followed by blocking with blocking buffer (5% non-fat dry milk, 50 mM Tris-HCl, 0.85% [w/v] NaCl, 0.05% [v/v] Tween-20, pH 7.5). The primary and secondary antibodies were applied in blocking buffer. The primary antibodies were mouse anti-FLAG (Sigma, F3165), mouse anti-HA-Tag (Santa Cruz Biotechnology, sc-7392), rabbit anti-CAPN5 (GeneTex, GTX103264), mouse anti-alpha 1 sodium-potassium ATPase (Abcam, ab7671), rabbit anti-ERK1 (Santa Cruz Biotechnology, sc-94), and mouse anti-β-actin (Sigma, A2228). The secondary antibodies were IRDye 800CW goat anti-rabbit IgG (Li-Cor, 926-32211) and Alexa Fluor 680 goat anti-mouse IgG (Thermo Fisher Scientific, A-21058). The images were acquired on the Li-Cor Odyssey CLx Imaging System.

### Confocal microscopy

The cells were seeded on gelatin-treated sterile coverglass (Zeiss, 474030-9000-000), and cultured as described above. One day later, the cells were fixed in 4% formaldehyde in 1× PBS (137 mM NaCl, 2.7 mM KCl, 10 mM Na_2_HPO_4_, 1.8 mM KH_2_PO_4_, pH 7.4), permeabilized with 1× PBS supplemented with 0.25% [v/v] Triton-X100, and blocked with 10% [w/v] BSA in 1× PBS. The antibodies were applied in 3% [w/v] BSA in 1× PBS. The primary antibody was rabbit anti-DDDDK Tag (Proteintech, 20543-1-AP), and the secondary antibody was Alexa Fluor 488 goat anti-rabbit IgG (Thermo Fisher Scientific, A-11034). The samples were treated with 4′,6-Diamidino-2-phenylindole (DAPI), mounted with ProLong Glass Antifade Mountant (Thermo Fisher Scientific, P36984), and imaged using a Nikon C2 confocal microscope with a 100× objective. Z-stack images of random viewfields were acquired with identical imaging parameters.

### Experimental rigor

All experiments were replicated a minimum of three times with representative results presented. Any adjustments in brightness/contrast were applied to the entire image. Blot lanes were not spliced from different images of a gel.

## Results

### CAPN5 is S-acylated on multiple cysteine residues

To determine whether CAPN5 was modified by S-acylation, we employed the acyl–PEG exchange (APEx) method [33]. The method is based on the replacement of fatty acyl groups attached via thioester linkage to cysteine residues with much larger polyethylene glycol (PEG) moieties. The attachment of the PEG groups is detected with denaturing gel electrophoresis followed by immunoblotting, where the number of shifted bands corresponds to the number of S-acylated cysteine residues in the protein.

C-terminal affinity tags were used to follow CAPN5 S-acylation. These detect full-length CAPN5 but not the autolytic fragments [2], and avoid complications of data interpretation resulting from the presence of autoproteolyzed CAPN5. Using C-terminally 3×FLAG-tagged WT CAPN5, we found that CAPN5 was S-acylated in both human neuroblastoma SH-SY5Y cells and human embryonic kidney HEK293T (293T) cells (Figure 1A). Four strong immunoreactive bands were detected, representing un-acylated CAPN5 (aligns with main band in negative control) and CAPN5 with 1, 2, or 3 PEG substitutions. A very faint fourth shifted band could also be observed. The number of shifted bands was highly reproducible in independent trials, with some variation in the intensities of the individual bands (Figure 1B).

**Figure 1.**
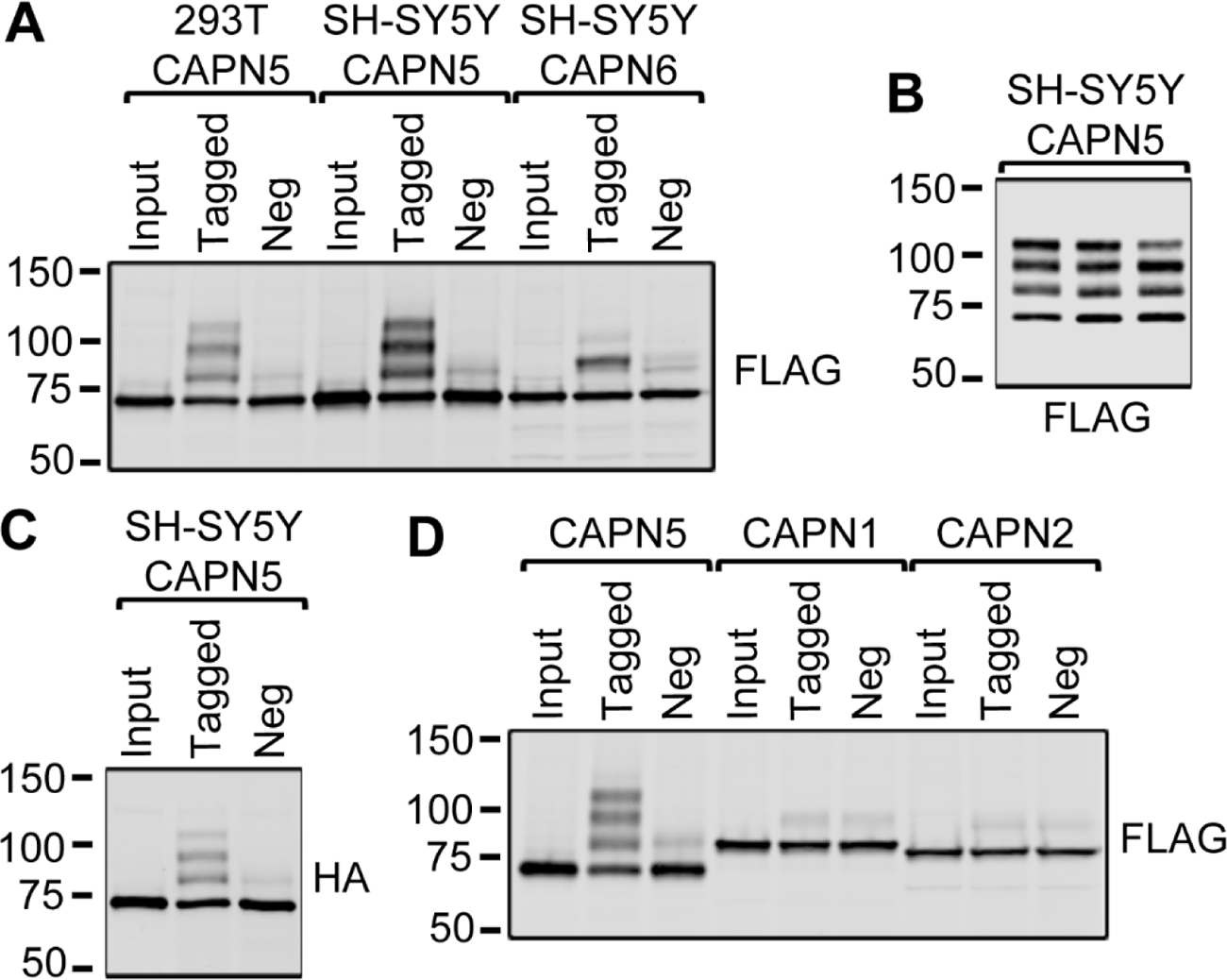
CAPN5 and CAPN6 are S-acylated, but CAPN1 and CAPN2 are not. Acyl-PEG exchange reactions were performed on total extracts of stable cells based on the human SH-SY5Y neuroblastoma cell line or the human 293T embryonic kidney-derived cell line, expressing 3×FLAG-tagged or 3×HA-tagged calpains, as indicated. The input, PEG-tagged (“Tagged”) and negative control (“Neg”) samples were resolved by denaturing gel electrophoresis, followed by immunoblotting with the indicated antibodies. **(A)** Both CAPN5 and CAPN6 were modified by S-acylation. Whereas CAPN5 molecules were modified at one to three residues, CAPN6 was modified at a single residue. **(B)** PEG-tagged samples from three independent acyl-PEG exchange reactions performed on total extracts of SH-SY5Y-based stable cells expressing WT CAPN5-3×FLAG, representing the reproducibility of the assay. **(C)** Acyl-PEG exchange reactions performed on total extracts of SH-SY5Y-based stable cells expressing WT CAPN5-3×HA, showing a similar modification pattern as with WT CAPN5-3×FLAG. **(D)** Acyl-PEG exchange reactions performed on total extracts of SH-SY5Y-based stable cells expressing the respective 3×FLAG-tagged calpains showed that unlike CAPN5, CAPN1 and CAPN2 were not modified by S-acylation. The molecular weights of the closest marker bands, represented by bars, are shown in kDa.

Similar results were obtained with C-terminally 3×HA-tagged CAPN5 in SH-SY5Y cells, suggesting that the S-acylation of CAPN5 was not dependent on the 3×FLAG affinity tag (Figure 1C). Quantification of the distribution of immunoreactive signal showed 41.4% of CAPN5 was unmodified and 58.6% ± 5.7% acylated from analysis of three independent trials. Negative controls consisted of parallel samples in which the thioester cleavage step was omitted. The single weak upshifted band observed in the negative controls suggests highly effective blocking prior to thioester cleavage with only a single cysteine resistant to blocking in a small population of CAPN5.

CAPN6 is the closest human paralog of CAPN5 and the only other known calpain with a C-terminal C2 domain. However, CAPN6 is catalytically inactive as it lacks the Cys residue in the active cite. Following stable expression of C-terminally 3×FLAG-tagged CAPN6 in SH-SY5Y cells, APEx revealed that CAPN6 is also S-acylated (Figure 1A). In contrast to CAPN5, CAPN6 was predominantly S-acylated at only a single site. To determine if S-acylation was unique to the CAPN5/6 subfamily of calpains, the catalytic subunits of classical calpains, CAPN1 and CAPN2, were expressed in SH-SY5Y cells and their S-acylation assessed using the APEx assay. S-acylation was not detected in CAPN1 or CAPN2 as the PEG-tagged and negative control lanes appeared identical (Figure 1D), although many reports suggest the transient association of calpains 1 and 2 with membranes or membrane proteins [37–40].

### Prediction of sites of S-acylation within CAPN5

Replacing individual cysteines with an alanine residue at potential sites of acylation is expected to selectively prevent acylation at a single site within a target protein. Destroying a single bona fide site of S-acylation by this tactic is expected to result in the loss of one of the upshifted immunoreactive bands resulting from the APEx chemistry. Because CAPN5 has 17 cysteines, S-acylation site prediction algorithms and examination of Cys residue conservation were used to help guide choices of residues for Cys-to-Ala mutagenesis.

S-acylation prediction algorithms used included CSS-Palm 2.0, NBA-Palm, MDD-Palm, PalmPred, and GPS-Lipid [41–45]. The results are summarized in Figure 2A and show that no cysteine residue was predicted by all algorithms to be an S-acylation site, but Cys-4, predicted by three out of five methods was most highly scored. Five additional Cys were predicted by two of the 5 methods and only 6 of the 17 cysteines, including the catalytic Cys-81 were not predicted by any method. The difficulty of predicting S-acylated cysteine residues in human proteins is due in part to the large number of protein S-acyltransferases in the human proteome [46, 47].

**Figure 2.**
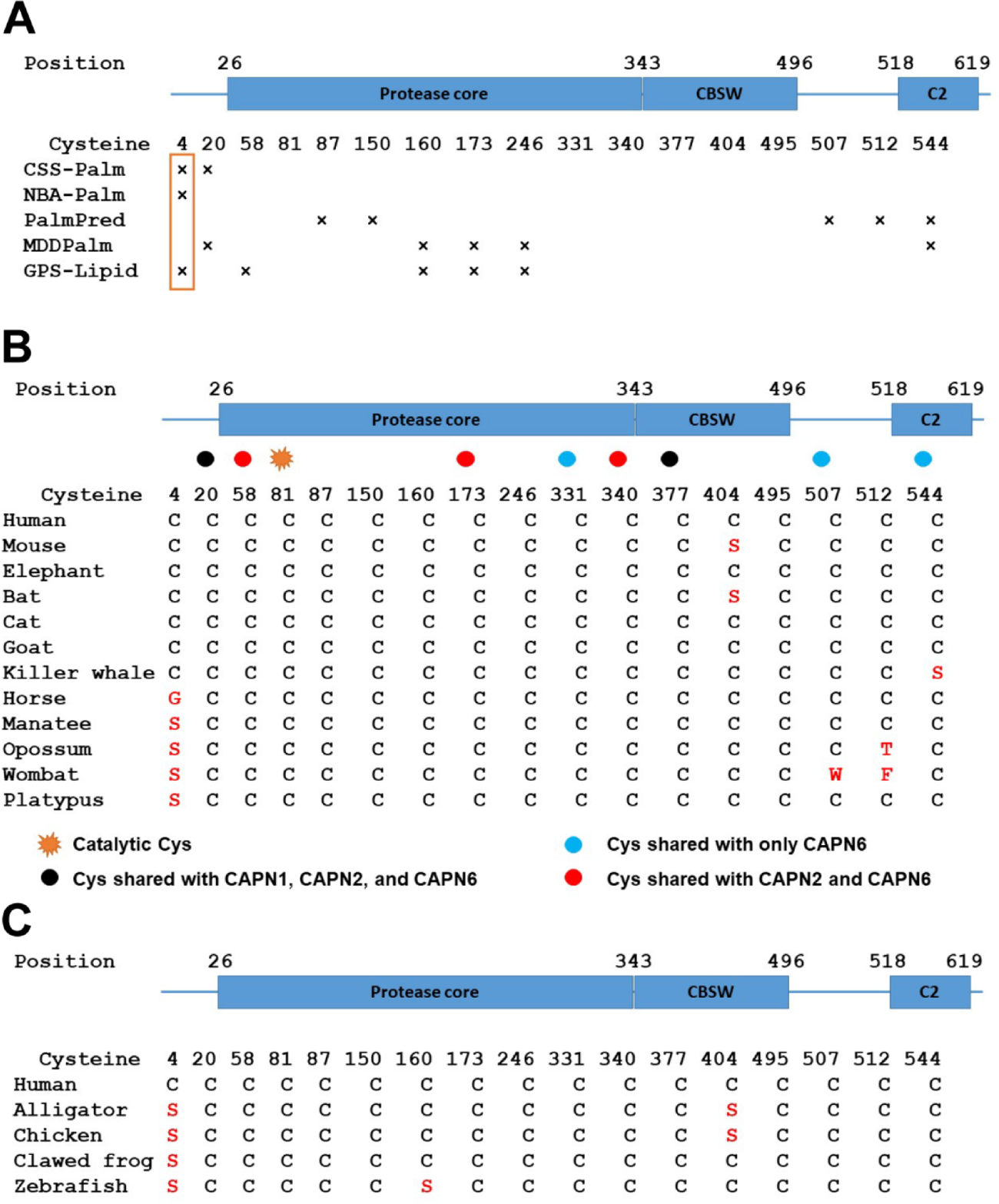
The prediction of S-acylated cysteine residues and the conservation of cysteine residues of human CAPN5. **(A)** The domain structure of CAPN5 (not drawn to scale) and the prediction of S-acylated cysteine residues in human CAPN5 by varied prediction algorithms. “×”, the respective cysteine position was predicted to be S-acylated. The most predicted residue was Cys-4 (boxed). **(B)** The conservation of the cysteine residues of human CAPN5 in Mammalia. The domain structure of CAPN5 (top) is not drawn to scale. The conservation of the cysteine residues of human CAPN5 in selected mammalian CAPN5 homologs is shown by “C”, and the substituted positions are marked by the respective one-letter code of the residue in red. The compared CAPN5 protein sequences were: human, *Homo sapiens* (order Primates) NP_004046; mouse, *Mus musculus* (order Rodentia) NP_001288179; African savanna elephant, *Loxodonta africana* (order Proboscidea) XP_003420073; greater horseshoe bat, *Rhinolophus ferrumequinum* (order Chiroptera) XP_032976748; domestic cat, *Felis catus* (order Carnivora) XP_003992755; goat, *Capra hircus* (order Artiodactyla) XP_017914797; killer whale, *Orcinus orca* (order Cetacea) XP_004279870; horse, *Equus caballus* (order Perissodactyla) XP_001494736; Florida manatee, *Trichechus manatus latirostris* (order Sirenia) XP_023592258; gray short-tailed opossum, *Monodelphis domestica* (order Didelphimorphia) XP_001377962; common wombat, *Vombatus ursinus* (order Diprotodontia) XP_027703985; and platypus, *Ornithorhynchus anatinus* (order Monotremata) XP_028904265. The cysteine residues of human CAPN5 shared with human CAPN1, CAPN2 and CAPN6 are shown with the respective indicated symbols. **(C)** The domain structure of CAPN5 (not drawn to scale) and the conservation of the cysteine residues of human CAPN5 as compared to selected vertebrate CAPN5 homologs. The conserved cysteine positions are marked by “C”, and the substituted positions are marked by the respective single-letter code of the residue in red. The compared CAPN5 protein sequences were: human, *Homo sapiens* NP_004046; alligator, *Alligator mississippiensis* XP_006264093; chicken, *Gallus gallus* XP_417278; clawed frog, *Xenopus laevis* NP_001080808; and zebrafish, *Danio rerio*. Please note that zebrafish has two CAPN5 isoforms, calpain-5A (NP_001073476) and calpain-5B (XP_001345114) with identical conservation pattern of the cysteine residues of human CAPN5. CBSW, calpain-type β-sandwich domain; C2, C2 domain.

Amino acid residues available for post-translational modifications are typically located in functional domains and a conserved sequence environment, as they confer some evolutionary constraints/stability on residues that are modified [48, 49]. We hypothesized that S-acylated cysteine residues in CAPN5 were likely conserved. A BLAST homology search [50] was performed against the RefSeq non-redundant protein database [51] and the positions of Cys in human CAPN5 are shown (Figure 2B). Conservation of these Cys positions within the closely related human CAPN6 and the ubiquitous isoforms CAPNs 1 and 2 were also examined (Figure 2B). Most cysteine residues of human CAPN5 were highly conserved in Mammalia (Figure 2B). Interestingly Cys-4, the most often predicted site for S-acylation, was conserved only among placental mammals, with the notable exceptions of substitution by glycine in the *Equus* (horse) genus and to serine in the manatee (*Trichechus manatus latirostris*). The Monotreme platypus (*Ornithorhynchus anatinus*) and the Marsupialians opossum (*Monodelphis domestica*) and wombat (*Vombatus ursinus*) also harbor a serine residue in the position that corresponds to human Cys-4 (Figure 2B). Among non-mammalian vertebrates, the position corresponding to Cys-4 in human CAPN5 was generally occupied by a serine residue (Figure 2C). Acylation and O-GlcNAcylation can occur at serine residues [52, 53], but was not evaluated in the present study. Only three Cys within CAPN5 were uniquely shared with CAPN6: at positions 331, 507 and 544. In addition to the catalytic Cys in CAPNs 1, 2 and 5, Cys at positions equivalent to 20 and 377 are conserved in all calpains examined here (CAPNs 1, 2, 5 and 6).

### Identification of sites of S-acylation within CAPN5

Despite Cys-4 being conserved only among placental mammals, this residue was predicted to be S-acylated by most algorithms (Figure 2A). Cys-20 is highly conserved in vertebrates and predicted to be S-acylated by two algorithms (Figure 2A and 2B). These two cysteine residues in the N-terminus of human CAPN5 are located similarly to the palmitoylated cysteine residues of the *P. falciparum* calpain [54], a unicellular protozoan parasite with a single calpain (Figure 3A). The N-termini of both proteins have clusters of positively charged amino acid residues (Figure 3A). Based on these similarities, we hypothesized that the Cys-4 and Cys-20 residues of human CAPN5 could be S-acylated. We observed that the Cys-to-Ala mutations altered the expression and/or turnover of CAPN5 (Supplemental Figure 1). Accordingly, we attempted to load similar levels of the un-modified CAPN5 for APEx analysis of S-acylation.

**Figure 3.**
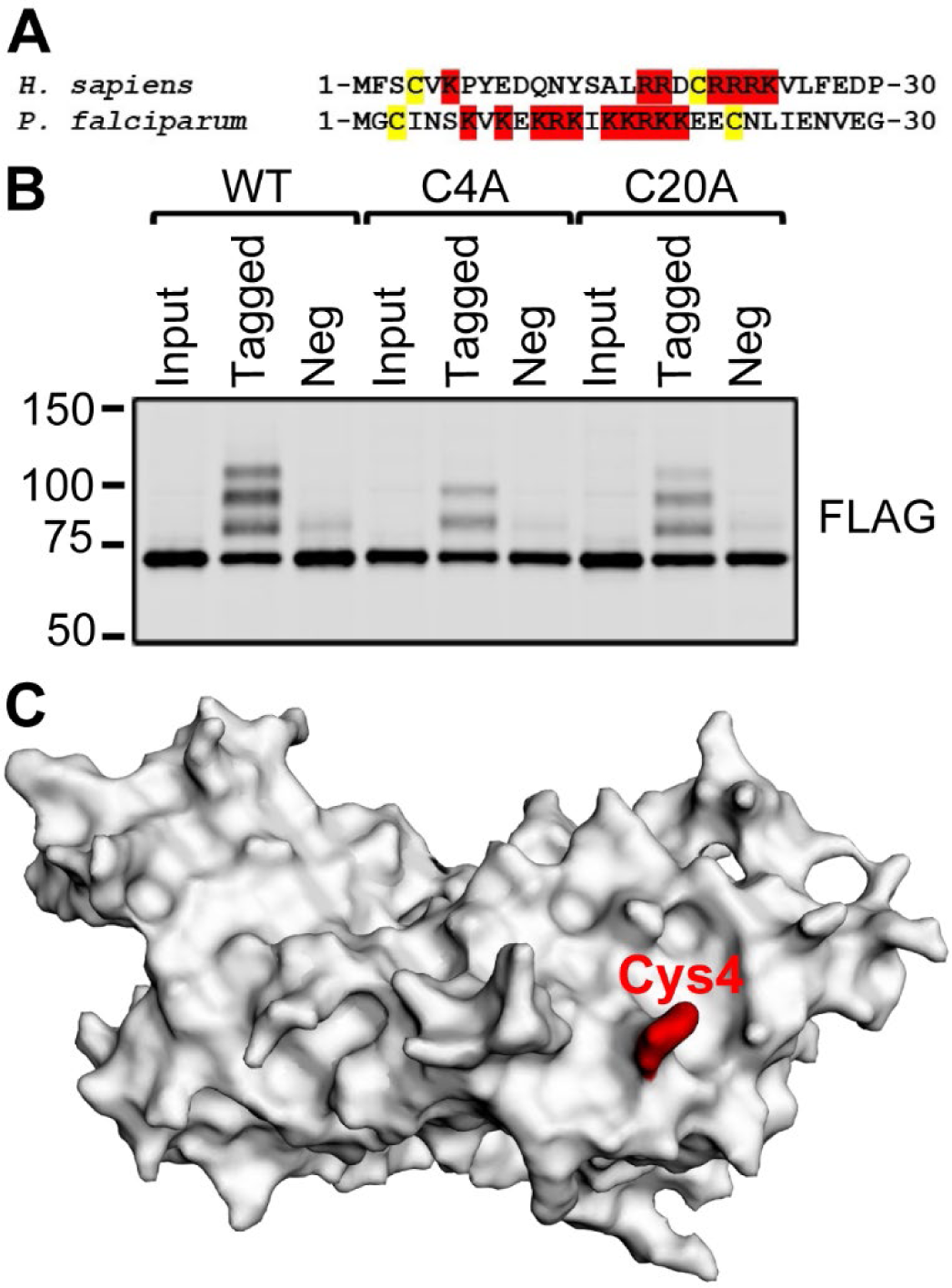
The Cys-4 residue of human CAPN5 is S-acylated. **(A)** Sequence alignment of the N-termini of human CAPN5 and *P. falciparum* calpain (NP_004046 and KNG77582, respectively). The cysteine residues are shown with a yellow highlight. Positively charged residues are shown with red highlight. **(B)** Acyl-PEG exchange reactions were performed on total extracts of SH-SY5Y-based stable cells expressing 3×FLAG-tagged CAPN5 constructs, as indicated. The input, PEG-tagged (“Tagged”) and negative control (“Neg”) samples were resolved by denaturing gel electrophoresis, followed by immunoblotting with anti-FLAG antibody. The loss of a single shifted band with the C4A mutant is consistent with the Cys-4 residue being S-acylated. However, the shifted band pattern of the C20A mutant was similar to that of WT CAPN5. The molecular weights of the closest marker bands, represented by bars, are shown in kDa. **(C)** The Cys-4 residue (shown in red) is exposed on the surface of CAPN5. Crystal structure of the CAPN5 protease core [55] visualized with EzMol [98].

Acyl-PEG exchange showed that the Cys-4 residue was S-acylated since the C4A mutant lost an upshifted band compared to WT CAPN5 (Figure 3B). In contrast, all three major upshifted bands were observed with C20A, although they were less intense than those associated with WT CAPN5 (Fig 3B). This may reflect less efficient acylation of the C20A mutant relative to WT CAPN5.

In an attempt to understand why the Cys-4 residue of human CAPN5 was S-acylated but Cys-20 was not, we inspected the crystal structure of the CAPN5 protease core domain [55] (Figure 3C). The CAPN5 core structure suggests that Cys-4 is surface exposed and thus would be accessible to an acyltransferase. However, Cys-20 was buried and inaccessible on the surface. S-acylation typically occurs on Cys residues exposed on the surface or located up to 8 Å into the inner leaflet of the membrane [56]. Further inspection of the CAPN5 core structure [55] suggested that Cys-246 was also partially exposed on the surface (Figure 4A), while remaining cysteine residues in the catalytic core domain other than Cys-4 were buried. Cys-246 is highly conserved and is located nearby His-252, a component of the Cys-His-Asn catalytic triad [5]. Acyl-PEG exchange revealed that S-acylation of the C246A mutant was severely impaired, however the three upshifted bands were faintly detected (Figure 4B). The detection of the three upshifted bands does not support Cys-246 and Cys-20 as a site of acylation.

**Figure 4.**
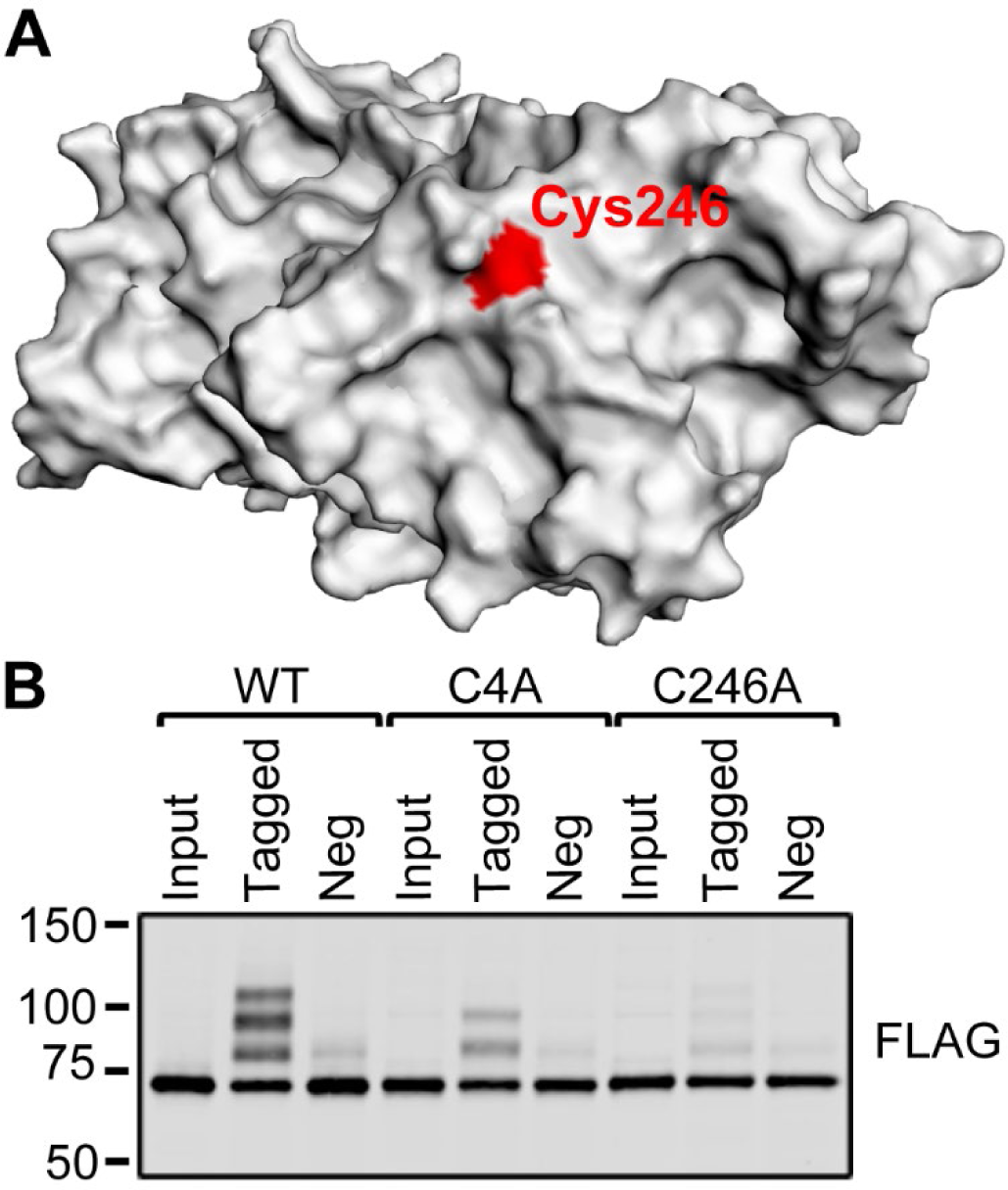
Mutating the Cys-246 residue severely impairs the S-acylation of CAPN5. **(A)** The Cys-246 residue is highlighted in red. Crystal structure of the CAPN5 protease core [55] visualized with EzMol. **(B)** Acyl-PEG exchange reactions were performed on total extracts of SH-SY5Y-based stable cells expressing 3×FLAG-tagged CAPN5 constructs, as indicated. The input, PEG-tagged (“Tagged”) and negative control (“Neg”) samples were resolved by denaturing gel electrophoresis, followed by immunoblotting with anti-FLAG antibody. The C246A mutation severely impaired the overall S-acylation of CAPN5, although a faint, WT CAPN5-like banding pattern was still detectable. The molecular weights of the closest marker bands, represented by bars, are shown in kDa.

Cys-377 is located within the calpain-type β-sandwich (CBSW) domain, also referred to as Domain III or C2-like domain [57]. Presently, there is no experimentally determined structure available for the CAPN5 CBSW domain. The Cys-377 residue was predicted to be partially surface exposed based on a homology model of the protease core-CBSW domain segment of CAPN5 generated by the Swiss-Model server (Figure 5A) [58, 59]. The other internal cysteine residue of the CBSW domain, Cys-404, was not predicted to be surface exposed. Although conserved between CAPNs 1, 2, 5 and 6, Cys-377 was not predicted to be an acylation site by any of the prediction algorithms. Acyl-PEG exchange showed that the overall S-acylation of the C377A mutant was severely impaired, but all three upshifted bands were faintly detectable (Figure 5B).

**Figure 5.**
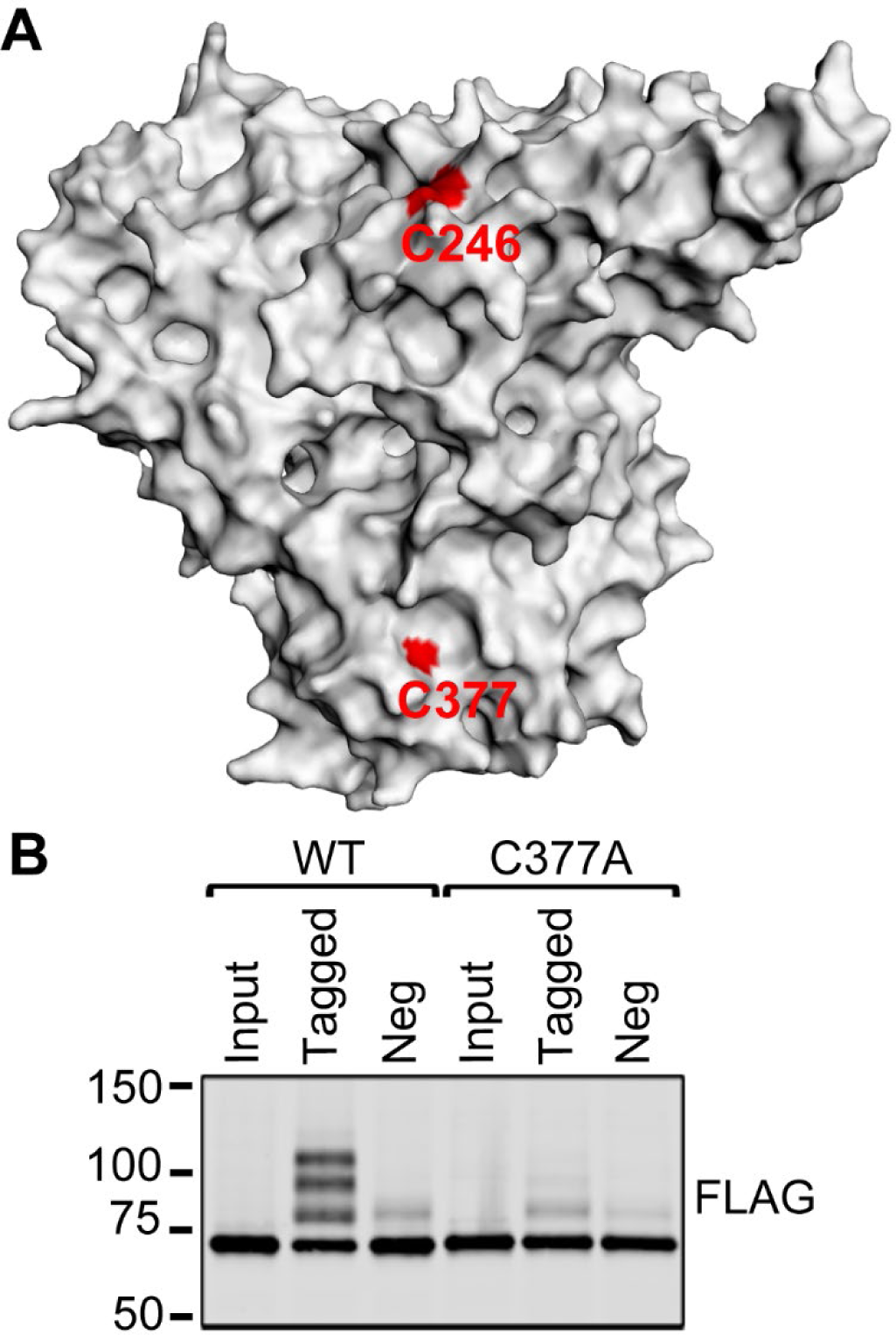
Mutating the Cys-377 residue severely impairs the S-acylation of CAPN5. **(A)** Swiss-Model homology model of the CAPN5 protease core – CBSW segment visualized with EzMol. The Cys-246 and Cys-377 residues are highligted in red. **(B)** Acyl-PEG exchange reactions were performed on total extracts of SH-SY5Y-based stable cells expressing 3×FLAG-tagged CAPN5 constructs, as indicated. The input, PEG-tagged (“Tagged”) and negative control (“Neg”) samples were resolved by denaturing gel electrophoresis, followed by immunoblotting with anti-FLAG antibody. The overall S-acylation of the C377A mutant was severely impaired, shown by the very faint shifted bands, as compared to WT CAPN5. The molecular weights of the closest marker bands, represented by bars, are shown in kDa.

The linker sequence between the CBSW and C2 domains of CAPN5 that harbors three Cys residues (Cys-495, Cys-507, and Cys-512) was also an attractive target to test for two reasons. One, results suggest that the CAPN5 C2 domain may position these proximal residues close to the membranes where the DHHC acyltransferases are anchored; and two, as a linker region, its modification is less likely to disrupt the overall folding/stability of the enzyme. The Cys-495 residue is at the C-terminus of the CBSW domain, and the Cys-507 and Cys-512 residues are between the CBSW domain and the C-terminal C2 domain (Figure 2).

Acyl-PEG exchange showed little S-acylation of the C495A/C507A/C512A triple mutant CAPN5 with almost no difference observed between the PEG-tagged and negative control lanes (total upshifted immunoreactivity = 5.51%, SD = 0.47%) (Figure 6A). In order to determine the contributions of the individual cysteine residues to the S-acylation of CAPN5, the single mutants C495A, C507A and C512A were evaluated.

**Figure 6.**
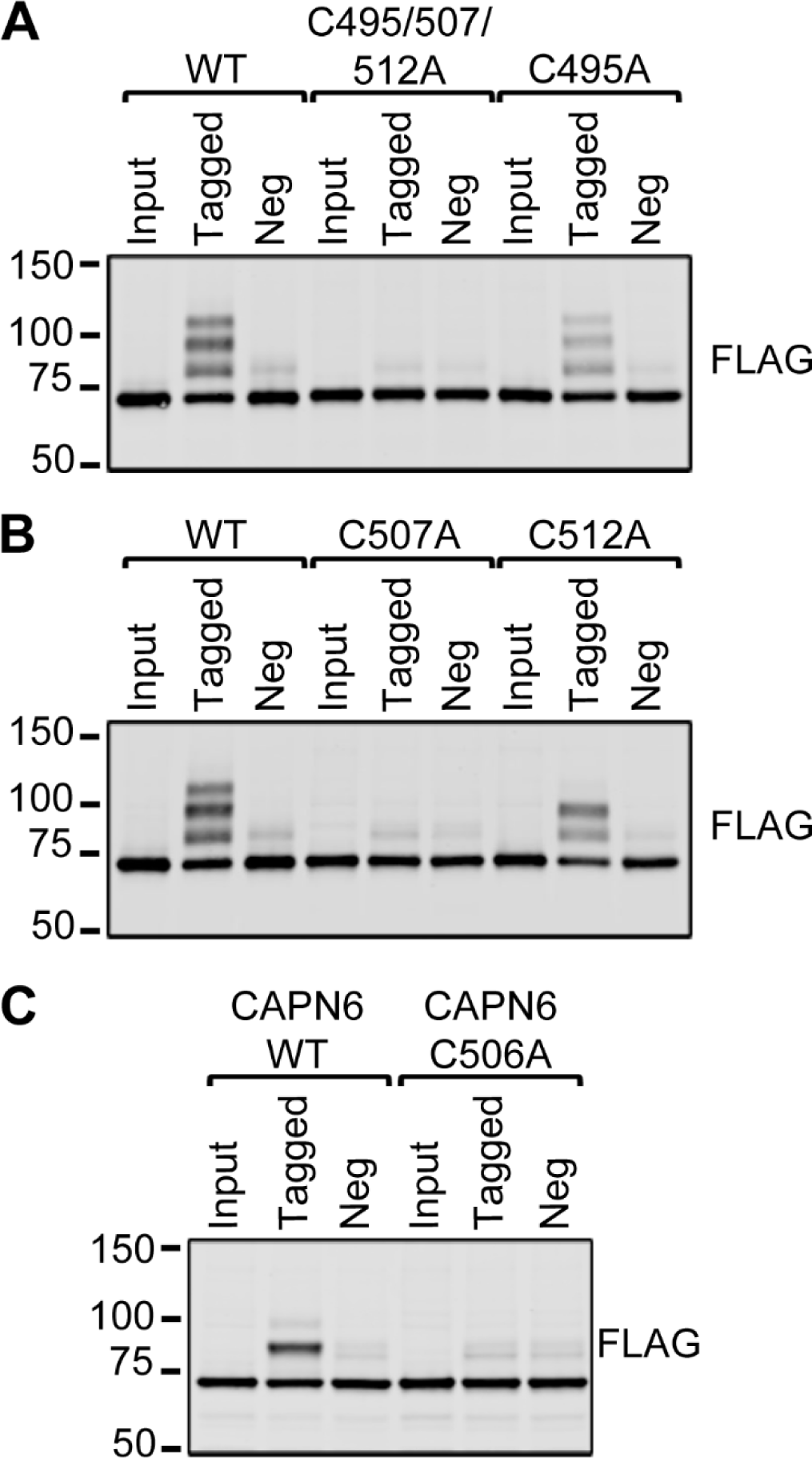
The role of Cys residues in the linker between the CBSW and C2 domains in S-acylation. Acyl-PEG exchange reactions were performed on total extracts of SH-SY5Y-based stable cells expressing 3×FLAG-tagged CAPN5 (A and B) or CAPN6 (C) constructs, as indicated. The input, PEG-tagged (“Tagged”) and negative control (“Neg”) samples were resolved by denaturing gel electrophoresis, followed by immunoblotting with anti-FLAG antibody. **(A)** The S-acylation of the C495A/C507A/C512A triple mutant CAPN5 was severely impaired. However, the C495A mutant CAPN5 showed a WT CAPN5-like banding pattern. **(B)** The S-acylation of the C507A mutant CAPN5 was severely impaired. The C512A mutant CAPN5 lost one shifted band as compared to WT CAPN5, consistently with the loss of a single S-acylation site. **(C)** The S-acylation of the C506A mutant CAPN6 was lost. The molecular weights of the closest marker bands, represented by bars, are shown in kDa.

For CAPN5-C512A there is a distinct loss of a single upshifted band, similar to what was observed for C4A. This identifies Cys-512 as a second site for S-acylation. In contrast, all three upshifted bands are distinctly visible for C495A leading to the conclusion that Cys-495 is not a site of S-acylation, although the intensity of the upshifted bands is diminished (total immunoreactivity shifted: 41.4%, SD = 2.61%). For C507A, there is little difference between the negative control and experimental (tagged) samples (total upshifted immunoreactivity = 9.65%, SD = 0.52%). This contrasts with C20A, C246A, C377A and C495A, where the three upshifted bands were detected, although decreased in intensity. The loss of upshifted bands with C507A suggests that Cys-507 may be a critical residue which must be acylated before Cys-512 and Cys-4 are able to be modified. The quantification of the APEx reactions of WT CAPN5 and its mutants is summarized in Figure 7.

**Figure 7.**
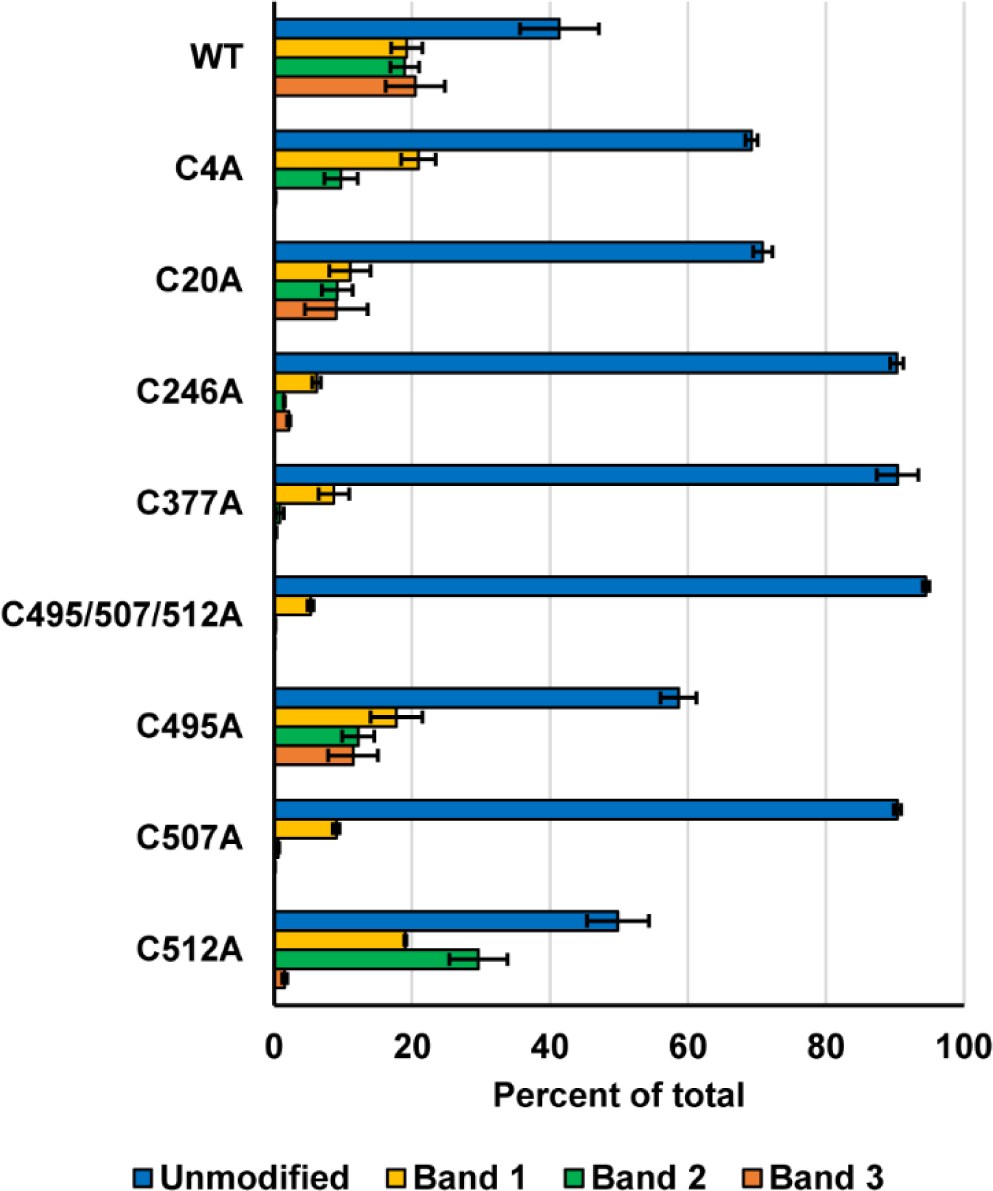
The quantification of the bands from acyl-PEG exchange reactions of WT and cysteine-to-alanine mutant CAPN5. The band intensities of the PEG-tagged samples from acyl-PEG exchange reactions of WT CAPN5 and its respective mutants were quantified from three independent trials, shown as percentages of their total intensities. Error bars, ± SD.

The Cys-4 and Cys-512 residues are not conserved in CAPN1, CAPN2 and CAPN6. However, the Cys-507 residue of CAPN5 corresponds to Cys-506 of CAPN6, whereas it is not conserved in CAPN1 or CAPN2 (Figure 2B). Accordingly, we hypothesized that the Cys-506 residue is the most likely site of the single S-acylation observed for CAPN6 (Figure 1A). Indeed, APEx analysis showed that the C506A CAPN6 mutant was not S-acylated (Figure 6C). This finding further supports that the Cys-507 residue of CAPN5 is directly involved in S-acylation. Inspection of the homology model of the CAPN5 C2 domain [2] suggested that the only cysteine residue within the C2 domain, Cys-544 is not exposed on the surface.

Recently a new structural model of full-length CAPN5, generated by the AlphaFold server [60, 61], became available. The AlphaFold CAPN5 model suggests that Cys-4, Cys-507 and Cys-512 are aligned near each other on the same side of the protein, and the side chains of all three residues are solvent accessible (Figure 8).

**Figure 8.**
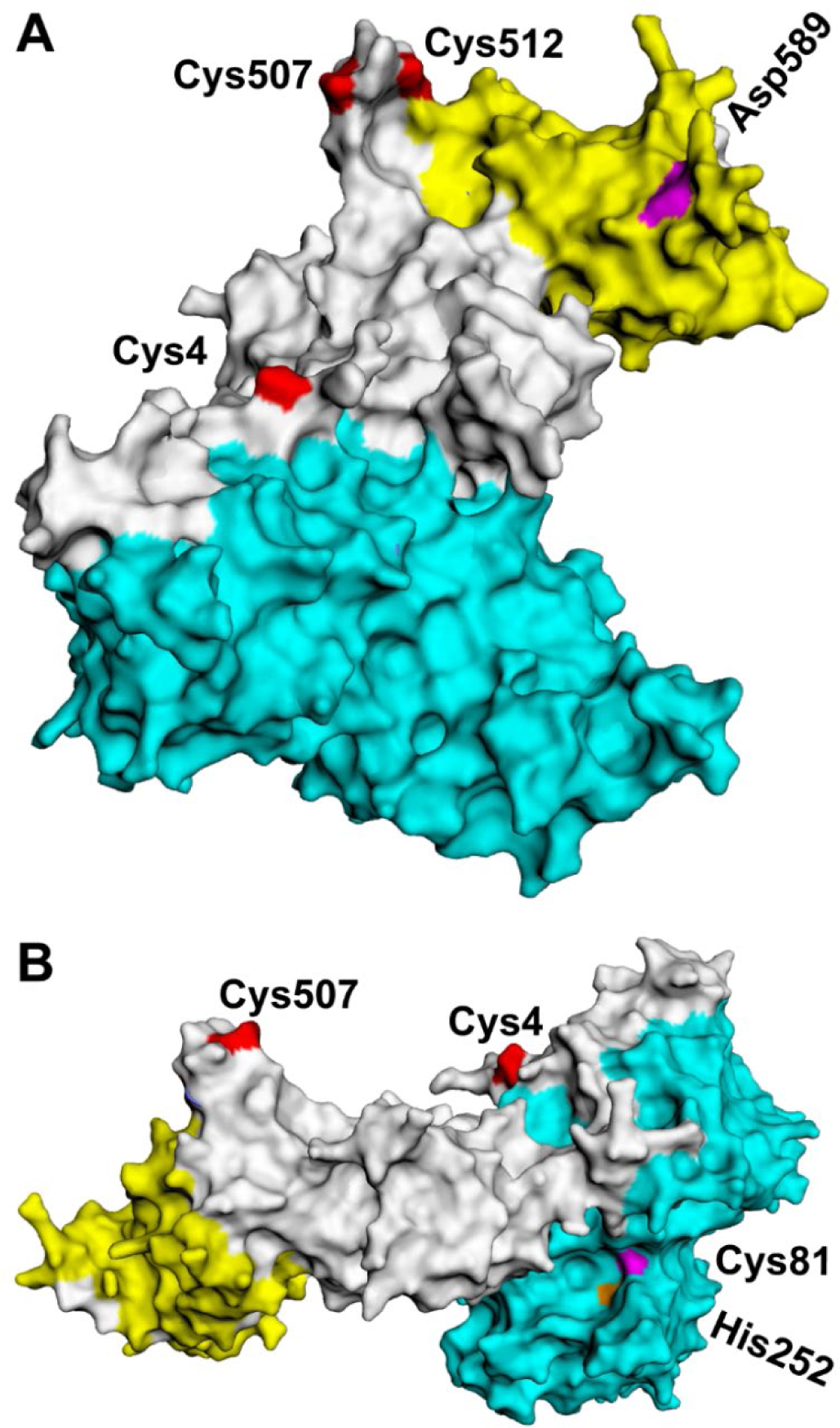
The AlphaFold model of CAPN5 shows close alignment of the S-acylated residues. **(A)** The Cys-4, Cys-507 and Cys-512 residues (highlighted in red) are aligned on the same side of CAPN5. Surface representation of the AlphaFold model of CAPN5, visualized with EzMol. The catalytic domain is highlighted in cyan, and the C2 domain is in yellow. The critical C2 domain residue Asp-589 is in magenta. **(B)** The side of CAPN5 reverse from that in (A) is shown with the catalytic residues Cys-81 and His-252 highlighted in magenta and orange, respectively.

### S-acylation is critical for the membrane association of CAPN5

Protein S-acyltransferases are intrinsic membrane proteins with active sites at the interface of the membrane and cytosol [62]. Proximity to those DHHC acyltransferases is required for S-acylation of CAPN5. CAPN5 and 6 are the only known calpains with C2 domains and having experimentally demonstrated here that each can be S-acylated, it seemed likely that the C2 domain could mediate their initial proximity to the membrane.

We recently documented that substituting asparagine for a highly conserved aspartate (D589N) within a potential calcium binding loop within the C2 domain of CAPN5 decreased its association with membranes and decreased the presence of the N-terminal fragment of CAPN5 produced by calcium dependent autoproteolysis [2]. If C2 domain-mediated membrane association is required for S-acylation, we predicted that the APEx analysis on cells expressing CAPN5-D589N-3×FLAG would show highly reduced S-acylation. For comparison lysates from cells expressing either the catalytically WT CAPN5 or the catalytically inactive variant CAPN5-C81A-3×FLAG were also analyzed (Figure 9A and B). For both the WT and C81A variants, more than half of the total CAPN5 immunoreactivity was upshifted (58.6%, SD = 5.7%; and 62.6%, SD = 3.2%, respectively) after the acyl-PEG exchange reaction. In contrast, the CAPN5-D589N, with no change to any cysteine residue, showed only very weak upshifted immunoreactivity (10.7%, SD = 3.1%) after the APEx reaction although there are three visible bands consistent with the presence of three modified cysteine residues. This suggests that the C2 domain has a role in localizing CAPN5 proximal to membrane associated DHHC acyltransferases, although other possible explanations for the overall loss of upshifted bands cannot be completely excluded at this time.

**Figure 9.**
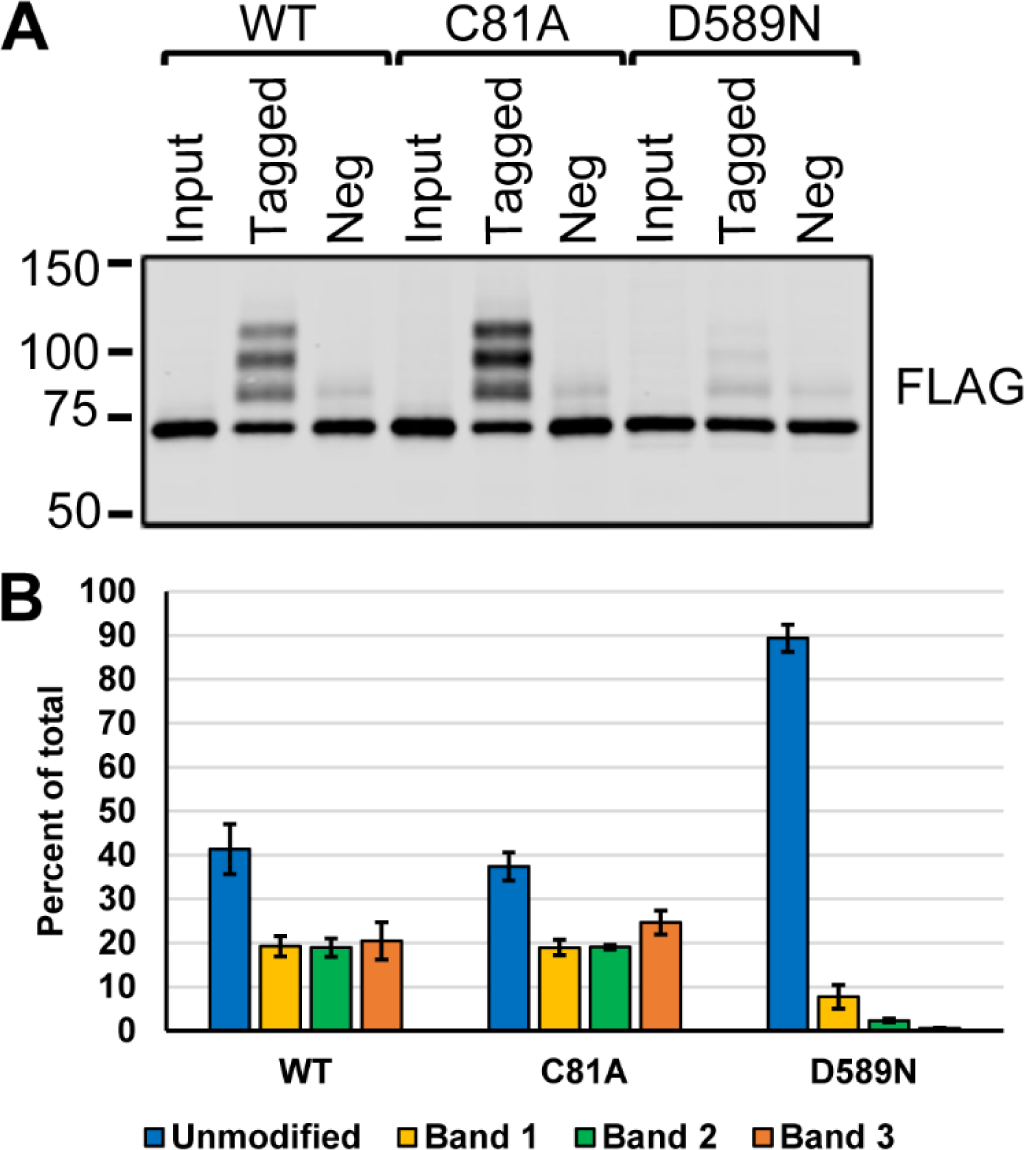
The intact C2 domain of CAPN5 is necessary for its efficient S-acylation, but its catalytic activity is not. **(A)** Acyl-PEG exchange reactions were performed on total extracts of SH-SY5Y-based stable cells expressing 3×FLAG-tagged CAPN5 constructs, as indicated. The input, PEG-tagged (“Tagged”) and negative control (“Neg”) samples were resolved by denaturing gel electrophoresis, followed by immunoblotting with anti-FLAG antibody. The S-acylation of the catalytically inactive C81A CAPN5 mutant was similar to that of the WT protein. However, the S-acylation of the C2 domain mutant D589N was severely impaired. The molecular weights of the closest marker bands, represented by bars, are shown in kDa. **(B)** The quantification of the band intensities of the PEG-tagged samples in **(A)** from three independent trials, shown as percentages of their total intensities. Error bars, ± SD.

Our current working model is that the C2 domain provides initial targeting of CAPN5 to the membrane for S-acylation of Cys-507, followed by acylation of Cys-512 and Cys-4 (Figure 10). We further hypothesized that S-acylation may help stabilize the membrane association of CAPN5. Confocal immunofluorescence experiments using anti-FLAG antibody show that WT CAPN5 localizes, at least in part, to plasma membrane ruffles and cilia (Figure 11). The C4A and C512A variants of CAPN5, missing Cys residues now shown to be specific sites of S-acylation, were still detectable in membrane ruffles and cilia but to a lesser extent than with WT CAPN5. Membrane association was dramatically reduced for cells expressing CAPN5 variants where the S-acylated fraction of the enzyme was shown to be <10% by APEx analysis (e.g., C246A, C377A and C507A).

**Figure 10.**
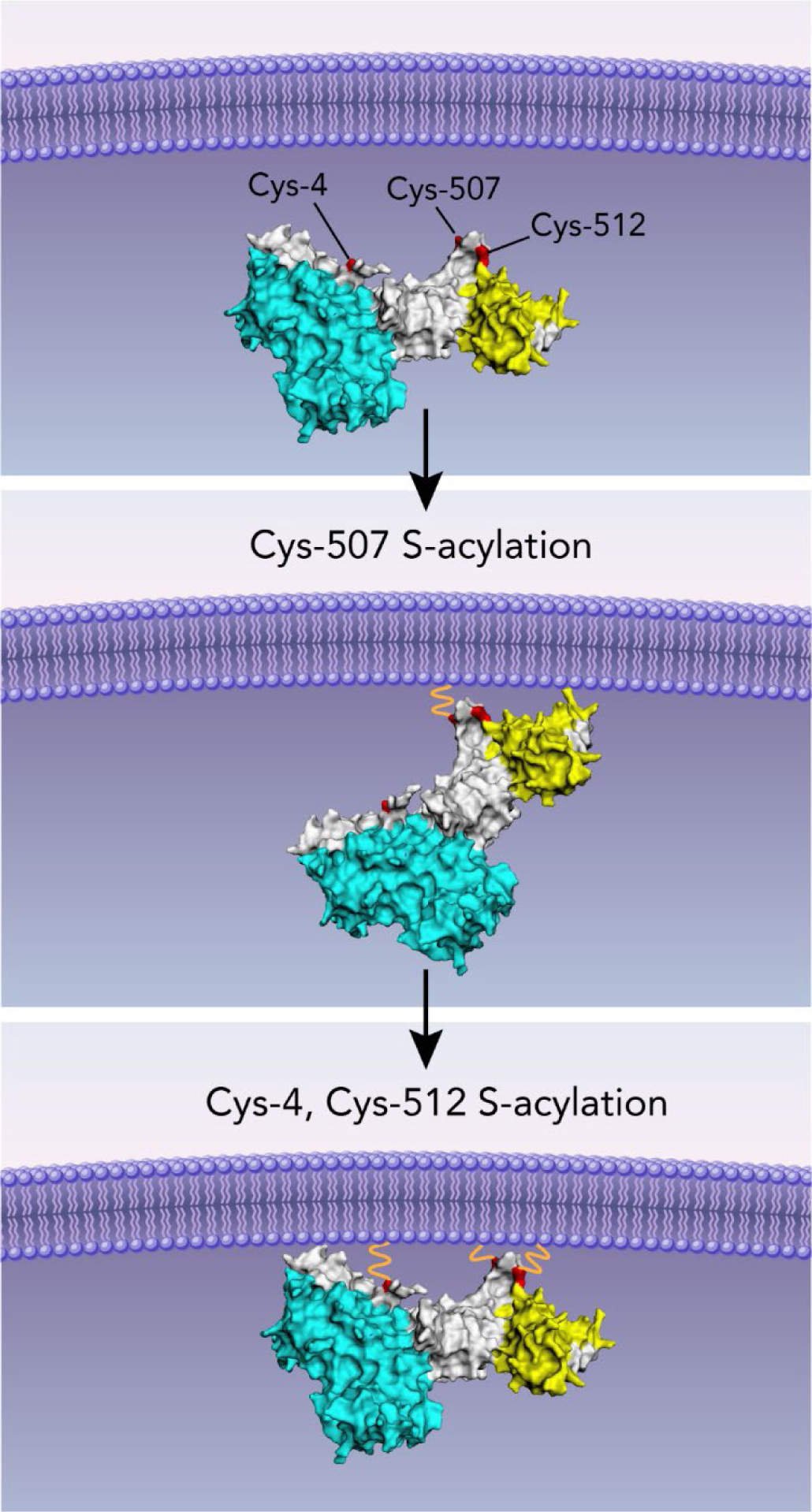
The structural model of CAPN5 offers clues to the function of S-acylation. Our proposed model showing the C2 domain-mediated membrane association of CAPN5, followed by the S-acylation of Cys-507, then that of Cys-4 and Cys-512. The S-acylation of CAPN5 positions its catalytic domain in close proximity to the membrane.

**Figure 11.**
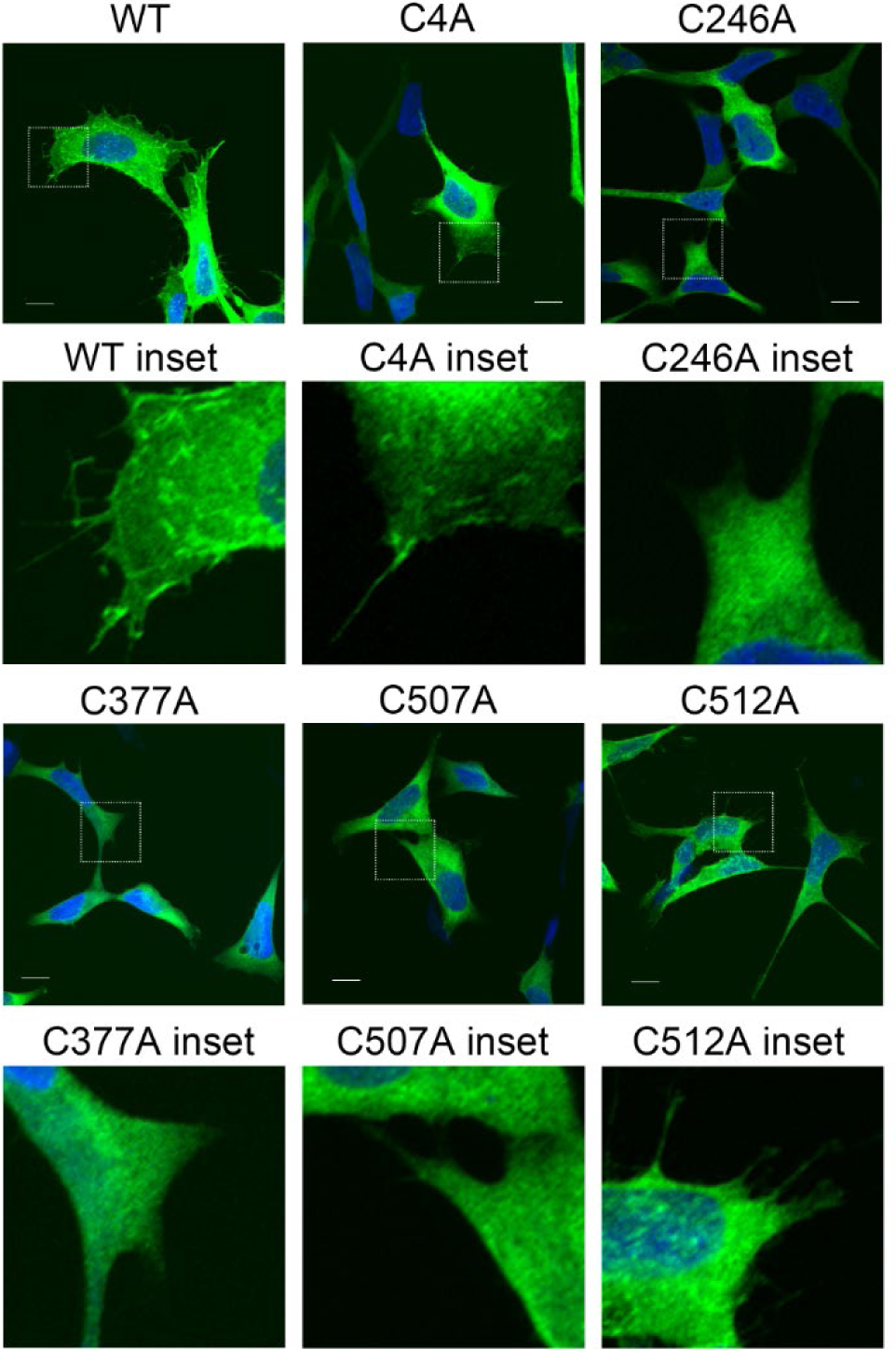
Decreased S-acylation interferes with the membrane association of CAPN5 in microscopy studies. Confocal microscopic images of anti-FLAG immunofluorescence of stable SH-SY5Y cells expressing the respective CAPN5-3×FLAG constructs. Wild-type CAPN5 was detected in membrane ruffles and cilia. This membrane-associated localization appeared weaker with the C4A and C512A mutants, but it was severely diminished with the C246A, C377A and C507A mutants, in good correlation with their S-acylation. Maximum intensity projections of Z-stacks are shown. The nuclei were stained with DAPI. Scale bars, 10µm.

Membrane association of CAPN5 was also evaluated by separation into cytosolic and membrane fractions, following incubation with CaCl_2_. We previously documented that membrane association favors CAPN5 activity in the presence of calcium as evidenced by the production of a predominantly membrane localized ∼ 61 kDa autoproteolyzed portion of CAPN5. This is detected using an anti-CAPN5 antibody, as the C-terminal 3×FLAG tag is within the region of CAPN5 removed by autoproteolysis [2]. Stable cells expressing the respective CAPN5-3×FLAG constructs were permeabilized and exposed to 2 mM calcium chloride, followed by separation into cytosol and membrane fractions, denaturing gel electrophoresis and immunoblotting to detect both full length and autoproteolyzed CAPN5. Consistent with earlier results [2], CAPN5-3×FLAG without altered Cys residues (WT) was partially membrane associated and the autoproteolytic fragment was predominantly present in the membrane fraction (Figure 12). Loss of either Cys residue (Cys-512 or Cys-4), determined to be sites of S-acylation, resulted in diminished detection of both full length CAPN5 and its autoproteolytic fragment in the membrane fraction. For the variants where total acylation was <10% by APEx experiments (C507A, C246A or C377A) there is diminished signal for full length CAPN5 in the membrane and for C246A and C377A, no visible autolytic fragment. There were faint signals of the autolytic fragment in the cytosolic fraction for WT CAPN5, C4A, and C507A. The fractionation studies showed that the loss of membrane association and autolysis of CAPN5 mutants was in good correlation with their degrees of S-acylation deficiency (Figure 7), and was also consistent with the immunofluorescent studies (Figure 11).

**Figure 12.**
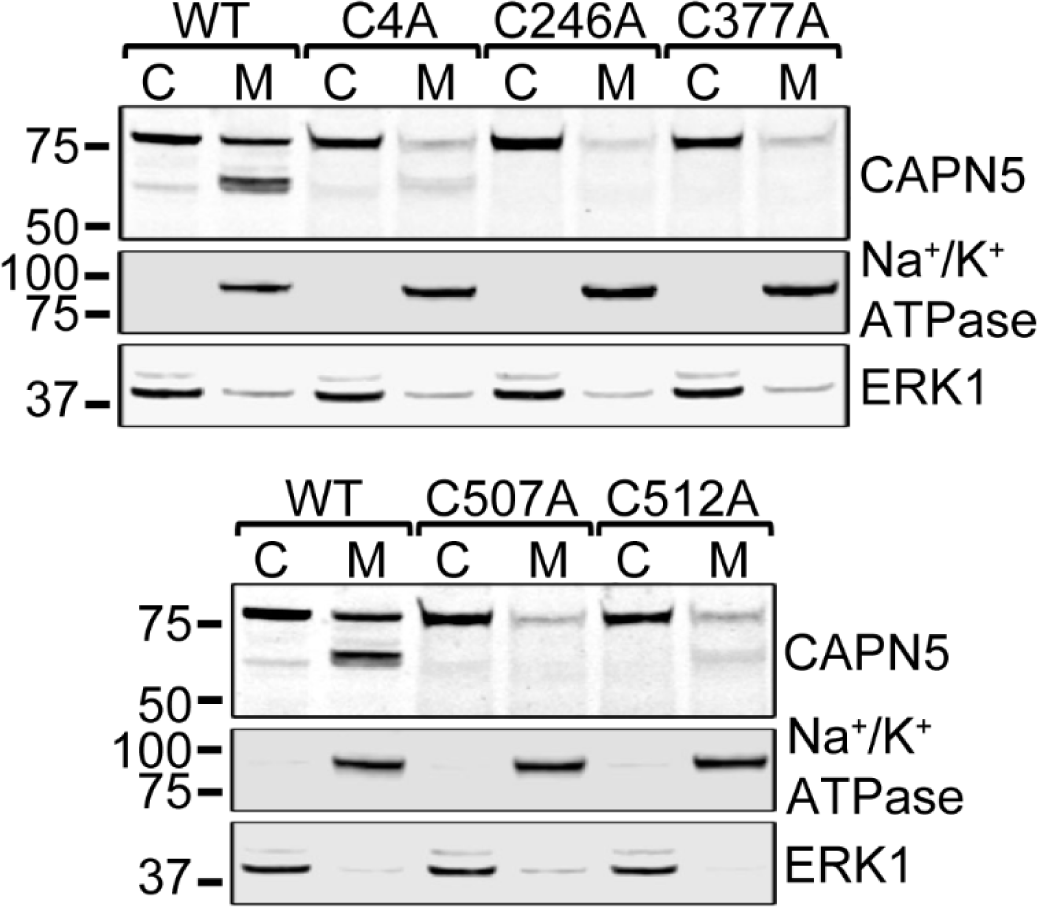
Decreased S-acylation interferes with the membrane association and activation of CAPN5 in fractionation studies. The respective SH-SY5Y-derived stable cells were permeabilized in the presence of 2 mM CaCl_2_, followed by the preparation of cytosolic (C) and membrane (M) fractions and denaturing gel electrophoresis. The proteins were transferred to nitrocellulose membranes, and probed with antibodies against CAPN5, sodium-potassium ATPase (membrane marker) and ERK1 (cytosolic marker). Wild-type CAPN5 was detected in both the cytosolic and membrane fractions. The autolytic N-terminal fragment of WT CAPN5 was detected primarily in the membrane fraction. The cysteine-to-alanine mutants whose S-acylation was impaired showed decreased membrane association and impaired autolysis. The molecular weights of the closest marker bands, represented by bars, are shown in kDa.

## Discussion

CAPN5 is present in both membrane and cytosolic fractions, and has been detected in the plasma membrane, nucleus, mitochondria, and synapses [14, 19–21, 63]. CAPN5 does not possess a transmembrane domain, but does contain a C2 domain which was recently confirmed to contribute to its membrane association [2]. Upon Ca^2+^-dependent activation, CAPN5 undergoes autolysis resulting in removal of part or the whole of the C2 domain [2]. Despite the loss of much of the C2 domain, the remaining N-terminal portion of calpain-5, which contains the catalytic core, remained associated with the membrane. This suggested that another mode of membrane association likely exists. We hypothesized that S-acylation/S-palmitoylation of CAPN5 might also play a role in its membrane association. The aim of this study was to evaluate the possible S-acylation of human CAPN5. Our results provide substantial evidence that CAPN5 is S-acylated. APEx reactions [33] with lysates of cells stably expressing CAPN5-3×FLAG, or a catalytically dead C81A variant, demonstrated that more than half of CAPN5 is modified with 1-3 acyl chains. S-acylation of CAPN5 was evident in both SH-SY5Y cells and HEK293T cells.

Supporting evidence that endogenously expressed CAPN5 can be S-acylated is found in large scale palmitoylomic studies. In the SwissPalm database, Release 3 (2019-09-08) [64], CAPN5 was identified in 4 out of 17 palmitoyl-proteomes of human cells [65–68], 4 of 17 mouse studies [69–72], and 1 of 2 rat studies [73]. CAPN5 was also identified in other studies describing the human palmitoylated proteome [74, 75]. None of these prior studies validated S-acylation in CAPN5 and did not specifically test for defined S-acylated Cys residues. Results from analysis of the APEx reactions reported here clearly demonstrate that CAPN5 is S-acylated on Cys residues 4 and 512. Corroborating evidence for the S-acylation of Cys-4 was found in results from a proteomic study of the human prostate cancer cell line DU145 that used acyl-biotin exchange (ABE) and affinity isolation of the biotinylated peptides [75]. The study identified an N-terminal peptide containing the Cys-4 residue of CAPN5. Importantly, the data document S-acylation at this site for endogenously expressed CAPN5. Interestingly, the unicellular protozoan parasite *Plasmodium falciparum* has only one calpain that was reported to be palmitoylated on both Cys-3 and Cys-22 [54]. However, the human CAPN5 APEx assays demonstrated that Cys-20 was not a site of S-acylation, suggesting that only a single N-terminal residue is similarly modified between *P. falciparum* and mammalian CAPN5. Although the Cys-4 residue is conserved in most placental mammals, other vertebrates have a serine residue in that position which doesn’t preclude the possibility of O-acylation in those species. Although not as extensively studied as S-acylation, O-acylation of serine residues can also occur such as that of Ser-3 in ghrelin [76].

An acyl-biotin exchange (ABE)-based S-acylation proteomics study using mouse forebrain tissue [71] identified three residues, Cys-495, Cys-507 and Cys-512 in the linker region between the CBSW domain (Domain III) and the C2 domain of CAPN5 as likely sites of S-acylation. However, due to the close proximity of these Cys residues, each of the sequence windows contained at least two potential S-acylation sites including Cys-507. Results reported here document that Cys-495 was not S-acylated, but that Cys-512 was modified as there was the expected loss of one upshifted immunoreactive band with the PEGylated C512A CAPN5-3×FLAG. For Cys-507 there was a more dramatic loss of two upshifted bands and only a small difference between the single upshifted band in the tagged sample above the negative control. Interpretation of the results for this site thus will be cautious. However, additional results suggest that Cys-507 is an important residue that must be S-acylated to facilitate the S-acylation of Cys-4 and Cys-512. The only other calpain with a designated C-terminal C2 domain is CAPN6, which was discovered at the same time as CAPN5, and is evolutionarily the closest relative of CAPN5 [12]. CAPN6 has a Lys residue instead of the active site Cys and is not proteolytically active. CAPN6 was also shown to be S-acylated, but was modified at only a single site, Cys-506 that corresponds to the Cys-507 residue of CAPN5. CAPN6 has not previously been identified in palmitoylomic studies, although this may be due to its restricted expression [12].

Membrane or lipid association has been reported for calpains-1 and 2, which also lack transmembrane domains [39, 77]. In the SwissPalm database, CAPN2 was found in 4/17 palmitoyl-proteomic studies for mouse cells [70, 78–80] and 1/17 for human [65], and CAPN1 was found only once in 17 studies for both species [65, 70]. However, APEx chemistry demonstrated that CAPN1 and CAPN2 are not S-acylated. A consensus mechanistic model called “kinetic trapping” has emerged to explain the S-acylation of soluble proteins. The long-chain fatty acyl groups are transferred to Cys residues by protein S-acyltransferases (PATs). All known mammalian PATs belong to the DHHC family of polytopic membrane proteins [27, 46, 47, 62, 81, 82]. According to the kinetic trapping model, S-acylation requires initial membrane association of soluble proteins mediated by either another lipid modification such as N-terminal myristoylation, C-terminal prenylation, or intrinsic membrane affinity of the target protein [83–87]. An acylated Cys residue is typically in close proximity to the lipid modification associated with the initial membrane attachment [31, 88]. For S-acylated transmembrane proteins, at least one acylated Cys is within 10 residues of the transmembrane domain boundary [32].

Our working model for S-acylation of CAPN5 is summarized in the schematic depicted in Figure 10. In this model, the C2 domain positions CAPN5 at the membrane followed by S-acylation of Cys-507, and then Cys-512 and/or Cys-4. For CAPN5, an intact C2 lipid association domain was necessary for its efficient S-acylation and two of the three acylated Cys residues were in close proximity to the C2 domain. The proposal that Cys-507 must first be modified to facilitate acylation of Cys-512 and/or Cys-4 is based on: 1) identification of the S-acylated peptide from endogenous CAPN5 [71], 2) it is one of the cysteines uniquely shared between CAPN5 and CAPN6, and the corresponding Cys-506 residue is the single site of S-acylation for CAPN6, and 3) its proximity to the C2 domain may favor its availability to one of the membrane-bound acyltransferases. There are other examples such as calnexin [89] and calcineurin [90] where S-acylation has been shown to be cooperative, as may be the case for CAPN5.

To the best of our knowledge, CAPN5 is the first protein whose S-acylation was shown to be facilitated by a C2 domain-mediated initial membrane association. To investigate if this mechanism is more widespread, we searched the UniProt/Swiss-Prot Knowledgebase (uniport.org) for proteins with C2 domains and obtained a list of 184 proteins from *H. sapiens*. Twenty-five of the 184 proteins with C2 domains were identified in one or more palmitoyl-proteomes in the SwissPalm database. Of these, ten were found in two or more of the 17 palmitoylomic studies: myoferlin, extended synaptotagmins 1 and 2, copine-3, calpain-5, neuropilin-1, protein kinase C alpha and gamma, 1-phosphatidylinositol 4,5-bisphosphate phosphodiesterase delta-3, and cyclin-G associated kinase. Synaptotagmin-1 was found in only one of seventeen palmitoyl-proteomes, however it is validated as being S-acylated [91, 92]. Thus, only a small subset of proteins with C2 domains appear to be also S-acylated.

Mutation of the Cys-246 and Cys-377 residues resulted in severe overall impairment of the S-acylation of CAPN5. We inspected the experimentally determined structure of the CAPN5 catalytic domain [55] and the AlphaFold model [60, 61] to investigate what could cause the impairment of S-acylation. Both the Cys-246 and Cys-377 residues are in an open coil segment where conversion to an alanine might be structurally disruptive, since alanine is the most favorable amino acid in an alpha helix [93]. Also, the side chain of Cys-377 is in a relatively polar environment where the alanine side chain would not be energetically favorable. Local structural disruption by either of these two mutations might therefore directly impair interaction with S-acylating enzymes or in some indirect manner prevent access to other acylation sites.

S-acylation can increase the stability of peripheral membrane proteins by protecting them from ubiquitination and degradation, target proteins for post-Golgi membranes, and regulate membrane trafficking and protein-protein interactions [31, 88, 94]. Altering specific Cys residues involved in CAPN5 S-acylation resulted in decreased cellular protein levels, consistent with decreased stability. Mutation of Cys-4 and Cys-512 decreased, but did not eliminate, the plasma membrane localization of CAPN5, whereas membrane localization was abolished by the C507A mutation. This is similar to Ras, which is localized to the Golgi or ER when non-palmitoylated, and to the plasma membrane when palmitoylated [86].

For CAPN5, a function of S-acylation appears to be the retention of the membrane association of the N-terminal portion including the protease core following Ca^2+^-dependent activation and autolysis which results in the loss of all or most of the C2 domain at the C-terminus. This contrasts with calpains 1 and 2 which undergo autolysis at the N-terminus, with autolyzed forms found exclusively in the cytosol [38, 95]. The N-terminal fragment of autolyzed CAPN5 contains the protease core and its retention at the membrane would presumably result in the cleavage of membrane-associated or membrane proteins. Several known calpain substrates are membrane-associated proteins [37, 40, 96, 97]. Prior to the discovery and characterization of CAPN5, the proteolysis of these substrates was often ascribed to the transient association of classical calpains, particularly calpain 2, with membrane phospholipids [39]. Our results suggest that at least some of the calpain-mediated proteolysis of membrane-proximal substrates is likely carried out by CAPN5. It will be important to determine the substrate spectrum of CAPN5 in future studies.

## Acknowledgments

We would like to thank our colleagues at the Spinal Cord and Brain Injury Research Center (SCoBIRC) and the Department of Neuroscience of the University of Kentucky for helpful discussions. We thank Matthew Hazzard, College of Medicine, University of Kentucky for the illustration in Figure 10. This work was supported by NIH grant R01 NS095229 (to James Geddes).

---

**Supplemental Figure 1.**
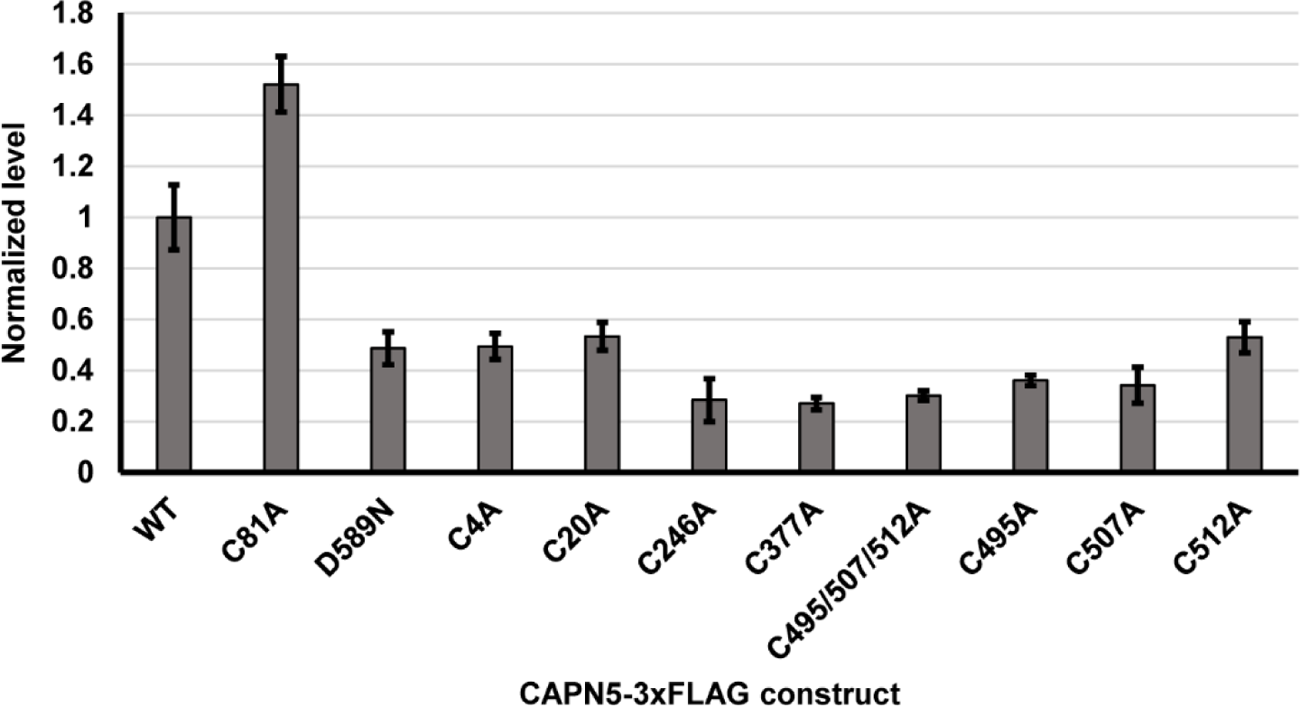
The relative levels of WT and mutant CAPN5-3×FLAG in the respective stable cells. The anti-FLAG immunoblotting band intensities representing WT or mutant CAPN5-3×FLAG were quantified in the input fractions of the acyl-PEG exchange reactions, and normalized to actin. The average normalized levels of three repetitions are shown, ± SD.

## Abbreviations used

3×FLAG: tag peptide with the sequence Asp-Tyr-Lys-Asp-His-Asp-Gly-Asp-Tyr-Lys-Asp-His-Asp-Ile-Asp-Tyr-Lys-Asp-Asp-Asp-Asp-Lys

3×HA: tag peptide with the sequence Tyr-Pro-Tyr-Asp-Val-Pro-Asp-Tyr-Ala-Gly-Ser-Tyr-Pro-Tyr-Asp-Val-Pro-Asp-Tyr-Ala-Gly-Ser-Tyr-Pro-Tyr-Asp-Val-Pro-Asp-Tyr-Ala

ABE: acyl-biotin exchange

ADNIV: Autosomal dominant neovascular inflammatory vitreoretinopathy

APEx: acyl-PEG exchange

APT: acyl-protein thioesterase

CBSW: calpain-type β-sandwich domain

PEG: polyethylene glycol

DAPI: 4’,6-diamidino-2-phenylindole

SD: standard deviation

FBS: fetal bovine serum

PEF: penta-EF hand domain

PAT: protein S-acyltransferase

PC: protease core domain

WT: wild-type

